# A checklist of alien taxa for South Africa

**DOI:** 10.1101/2025.05.22.655507

**Authors:** TA Zengeya, KT Faulkner, MP Mtileni, L Fernandez Winzer, S Kumschick, EJ McCulloch-Jones, SA Miza-Tshangana, TB Robinson, A Sifuba, W Engelbrecht, BW van Wilgen, JRU Wilson

## Abstract

This paper presents what is intended to be a comprehensive checklist on alien taxa in South Africa developed as part of triennial national status reports on biological invasions. It thus includes: taxa that are, have been, or were proposed to be regulated; alien taxa that are or have been present in South Africa (including those only ever recorded in quarantine facilities); taxa that are native to a part of South Africa that have formed native-alien populations in another part of the country; and taxa which have been recorded at some point as alien or for which the risk of invasion has been evaluated. Names used previously are included so it is clear why taxa listed in historical sources are no longer considered alien or present, and how such names have been interpreted in terms of the latest authoritative taxonomic sources. The list also includes information on the invasion status of the taxa, their pathways, distributions, impacts, and management, with metadata provided for all 38 variables, including confidence and data sources for 23 of them. The development of documented and repeatable workflows ensures it is clear why taxa (and associated information) are included on the list and facilitates reviews and updates. Based on information up to the end of December 2022, the checklist includes over 6000 taxa, of which over 3500 are alien taxa confirmed as present outside of captivity or cultivation. However, several key data sources still need to be verified and integrated into the list (particularly taxa in captivity or cultivation). Thus, this list should not yet be regarded as a complete baseline of the knowledge of alien taxa present in South Africa. The checklist is presented in a manner that is tidy and FAIR (findable, accessible, interoperable, reusable) and will be maintained, expanded, and updated, with the aim for the list to become comprehensive and dynamic. By so doing, the checklist will allow the number and status of alien taxa to be tracked over time, informing management planning and regulatory decisions.

## Background and summary

South Africa is a large, mega-biodiverse country that has a long history of alien taxa introductions and invasions (Faulkner et al. 2020, van Wilgen et al. 2020a). Alien taxa in South Africa come from a range of taxonomic groups, occur in a variety of habitats, and have had pervasive impacts (van Wilgen et al. 2022b). In an effort to improve our knowledge on biological invasions in South Africa, and to inform invasive species management and policy, various lists of alien taxa have been developed (Faulkner et al. 2015). These lists each have their own purpose and focus. For example, taxon-, habitat-, and site-specific lists of aliens have been compiled for freshwater fishes (Ellender and Weyl 2014), marine organisms (Robinson et al. 2020a), terrestrial molluscs (Herbert 2010), and the Prince Edward Islands (Fernández Winzer et al. 2024, 2025).

Lists of alien taxa are challenging to produce and maintain (McGeoch et al. 2012), not least because they are dynamic, with new taxa being added to the lists as they are introduced and/or detected, and others being removed as they die out or are eradicated (Matthys et al. this issue, van Wilgen et al. 2020a). However, many South African lists are static and are produced as one-off publications [though updated versions of lists of alien marine organisms (Robinson et al. 2020a) and biological control agents (Zachariades 2021) are published periodically]. Lists also differ between curators. For example, the numbers reported for alien terrestrial vertebrates in South Africa vary greatly across lists (van Wilgen et al. 2020a). These discrepancies are partly due to the lists being compiled using different methods, but also because different lists implement different standards and definitions (van Wilgen et al. 2020a).

Lists of alien taxa often provide important ancillary information on, for example, introduction pathways, distributions, invasion status, and dates of first record (Faulkner et al. 2015). However, as for the checklists themselves, there is significant variation in what ancillary information is presented and in the completeness of this information (Faulkner et al. 2015), and sources and confidence levels are rarely consistently and systematically included. Global standards have been developed for alien species data that are maintained by the Darwin Core team (Groom et al. 2019), but, to date, South African lists of alien taxa have not followed these standards or have not explicitly tried to meet FAIR data principles (Findable, Accessible, Interoperable, and Reuseable; Wilkinson et al. 2016).

A comprehensive, up-to-date list of alien taxa in South Africa is required to provide estimates on the status and trends of biological invasions, and to get an idea of whether interventions are effective. In particular, such estimates are required for South Africa’s triennial report ‘The status of biological invasions and their management in South Africa’, which the South African National Biodiversity Institute is mandated to produce under the Alien and Invasive Species Regulations of the National Environmental Management: Biodiversity Act (NEM:BA A&IS Regulations; Department of Environment, Forestry and Fisheries, 2020). Three reports have been produced to date, each with an accompanying list (van Wilgen and Wilson 2018, Zengeya and Wilson 2020, 2023a). A list of alien taxa is also required for South Africa to measure and report on its progress towards global conservation targets, such as Target 6 of the Kunming-Montreal Global Biodiversity Framework.

Building on the list produced for the last national status report (SANBI and CIB, 2023a), here we present a consolidated checklist of alien taxa for South Africa that provides current knowledge on their status, and information on their pathways, distributions, impacts, and management. This checklist follows global biodiversity data standards, and the data are intended to be tidy and FAIR. The intention is also for this list to form a baseline for updating other relevant lists [e.g., the Global Register of Introduced and Invasive Species for South Africa that lists invasive taxa that are recorded (or in this case presumed) to have negative impacts (Robinson et al. 2020b)].

## Information about the data

### Region

Mainland South Africa and inshore islands [a separate list is curated for the Prince Edward Islands, South Africa’s sub-Antarctic territories (SANBI and CIB 2023b; Fernández Winzer et al. 2024, 2025)].

### Period of study

Sources published or available up to 31 December 2022, with sources dating from 1906 (Theobald 1906). Some alien taxa were introduced to South Africa prior to European colonisation in the second half of the 17^th^ century, but the majority were introduced since then (Faulkner et al. 2020).

### Objective

To compile a consolidated checklist of alien taxa in South Africa updated in-line with triennial national status reports, with the aim, in future, to update as information becomes available (with information available through a dashboard; see Zengeya et al. this issue for more details).

### Source of funding

This project was funded by the Department of Forestry, Fisheries and the Environment (DFFE) through the South African National Biodiversity Institute (SANBI) and the DSI-NRF Centre of Excellence for Invasion Biology (CIB).

### Methodology

The checklist of alien taxa in South Africa is the result of a process to consolidate and standardise information on the presence of alien taxa in South Africa from various sources. The checklist is intended to primarily record the presence of alien taxa in South Africa (be they inside or outside of captivity or cultivation). However the list also includes some taxa that are not present in the country, in particular, those that were listed as prohibited under the NEM:BA A&IS Regulations of 2014 or 2016 or in draft lists of the regulations in 2007, 2009, 2013, 2014, 2015, and 2016 (prohibited taxa are those that were believed to be absent but posed a significant risk of invasion; Wilson and Kumschick 2024). Moreover, some taxa which are native to South Africa are also included: in particular taxa with native-alien populations (*sensu* Nelufule et al. 2022), and taxa that are included as mandated by the NEM:BA A&IS Regulations of 2020 [e.g., ‘indigenous’ taxa for which risk analyses have been completed; for a full discussion see ‘isNative’ in the metadata (SANBI and CIB 2023c)].

Data were extracted from various sources and merged based on standardised taxonomy (Figure 1). The processes followed are documented and described below.

**Figure 1.**
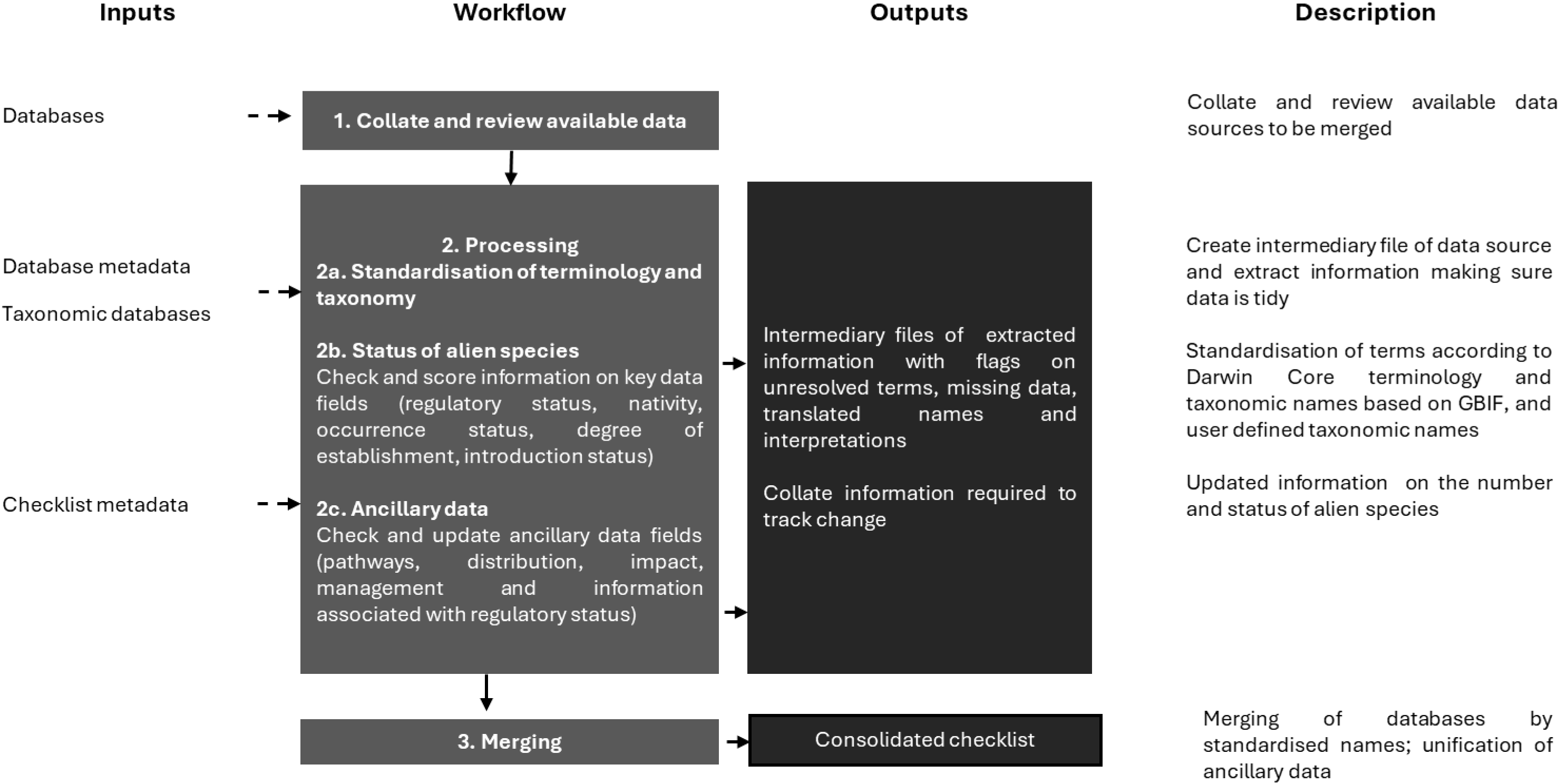
Overview of the workflow that was used to compile a consolidated checklist of alien taxa for South Africa

### 1. Collation and review of available data

The data sources varied (e.g., government reports, peer-reviewed papers, grey-literature, atlassing projects and online databases) and contained various types of information (Table 1). Most data sources were peer reviewed literature and included inventories of plants (e.g., Glen 2002; Hoy et al. 2021), microbes (e.g., Wood 2017; Paap et al. 2018), invertebrates (e.g., Prinsloo and Uys 2015; Hurley et al. 2017; Janion-Scheepers et al. 2020), amphibians (e.g., Measey et al. 2020), fishes (e.g., Ellender and Weyl 2014; Weyl et al. 2020), reptiles (e.g., van Rensburg et al. 2011; Measey et al. 2020), birds (e.g., Macdonald et al. 1986; Picker and Griffiths 2017), mammals (e.g., van Rensburg et al. 2011, Measey et al. 2020). Several sources contained context-specific inventories of alien taxa, for example, aquatic animals (De Moor and Bruton 1988), those in the pet trade (e.g., Nelufule et al. 2020), protected areas (Foxcroft et al. 2023), marine taxa (Robinson et al., 2020a), and the regulatory lists (Wilson and Kumschick 2024). Information was also obtained from databases such as the Barcode of Life Data System (BOLD; http// www.barcodinglife.org), Southern African Bird Atlas Project 2 (SABAP2) (http://sabap2.birdmap.africa/), the Botanical Database of Southern Africa (BODATSA) (http://posa.sanbi.org/); and the Southern African Plant Invaders Atlas (SAPIA).

**Table 1.**
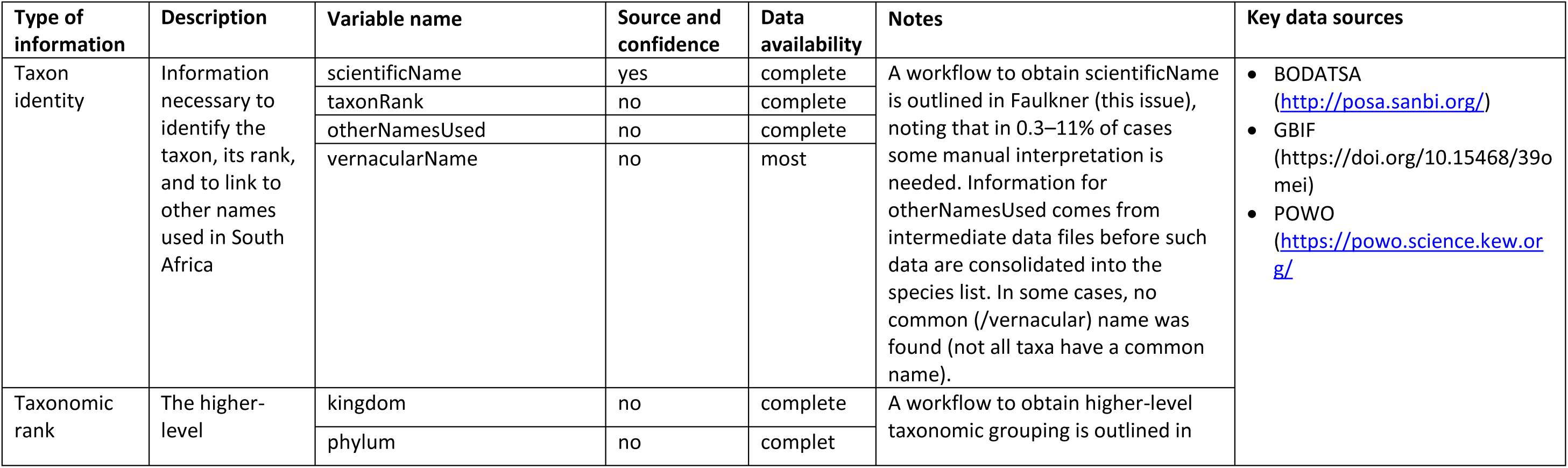
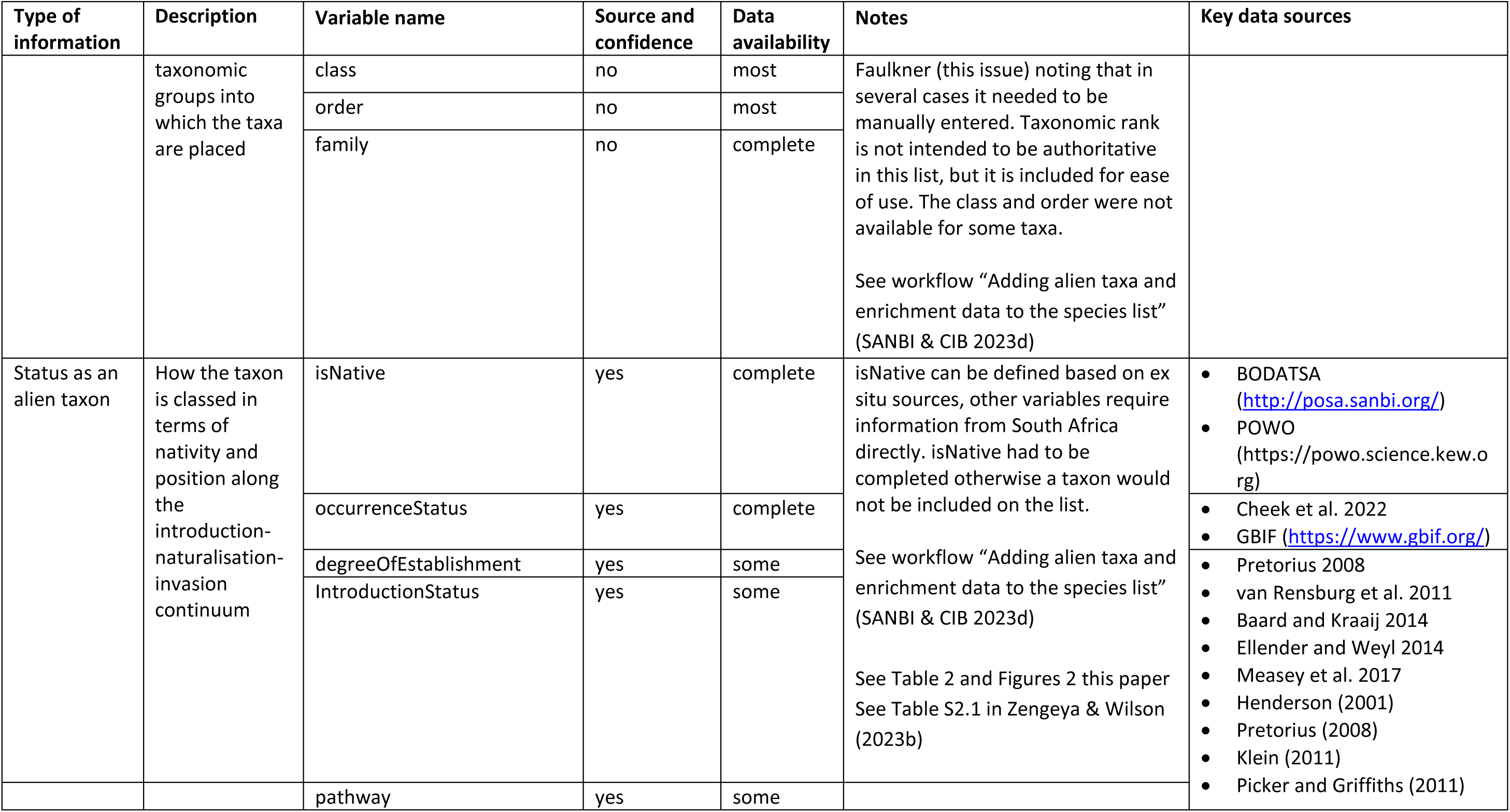
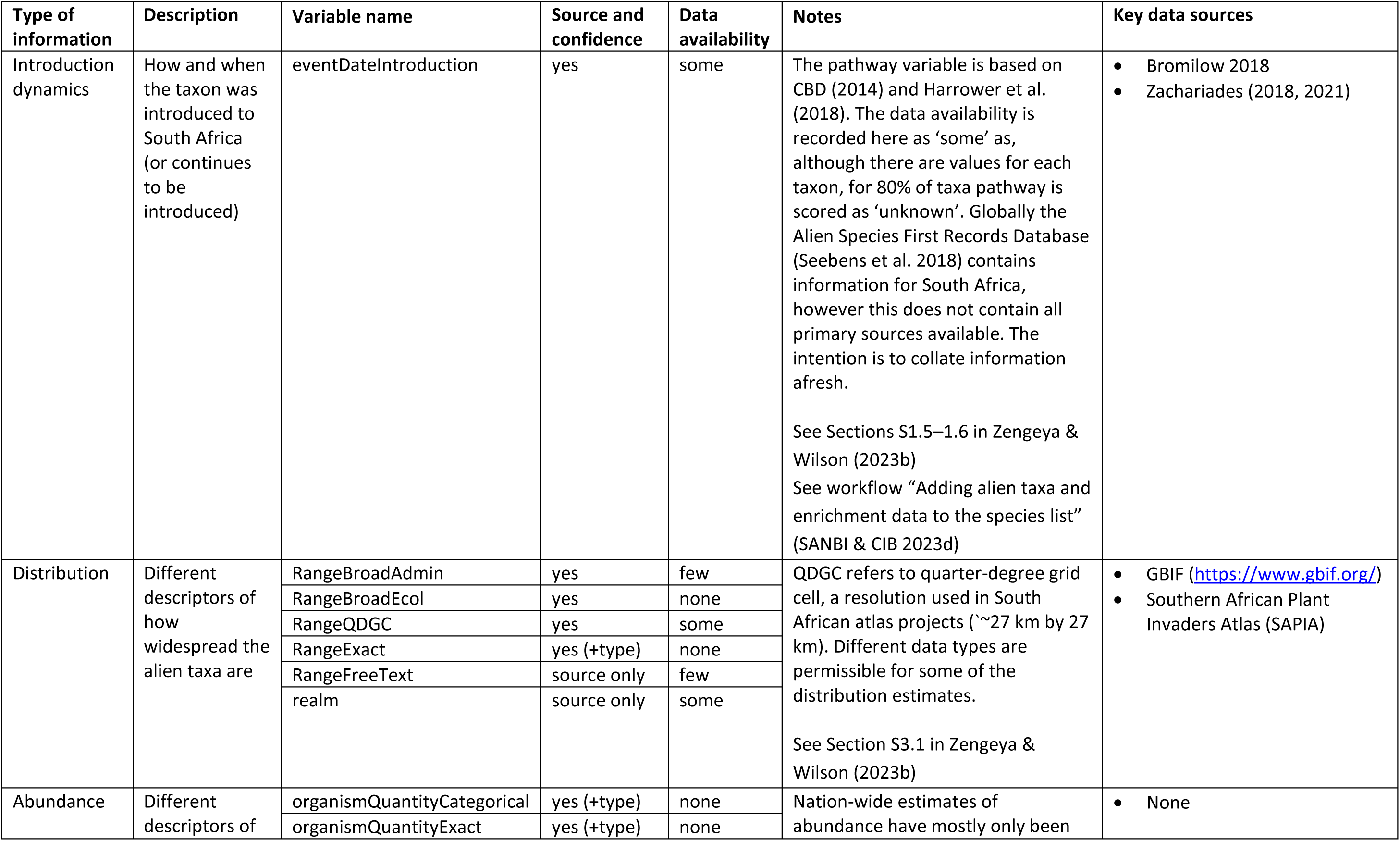
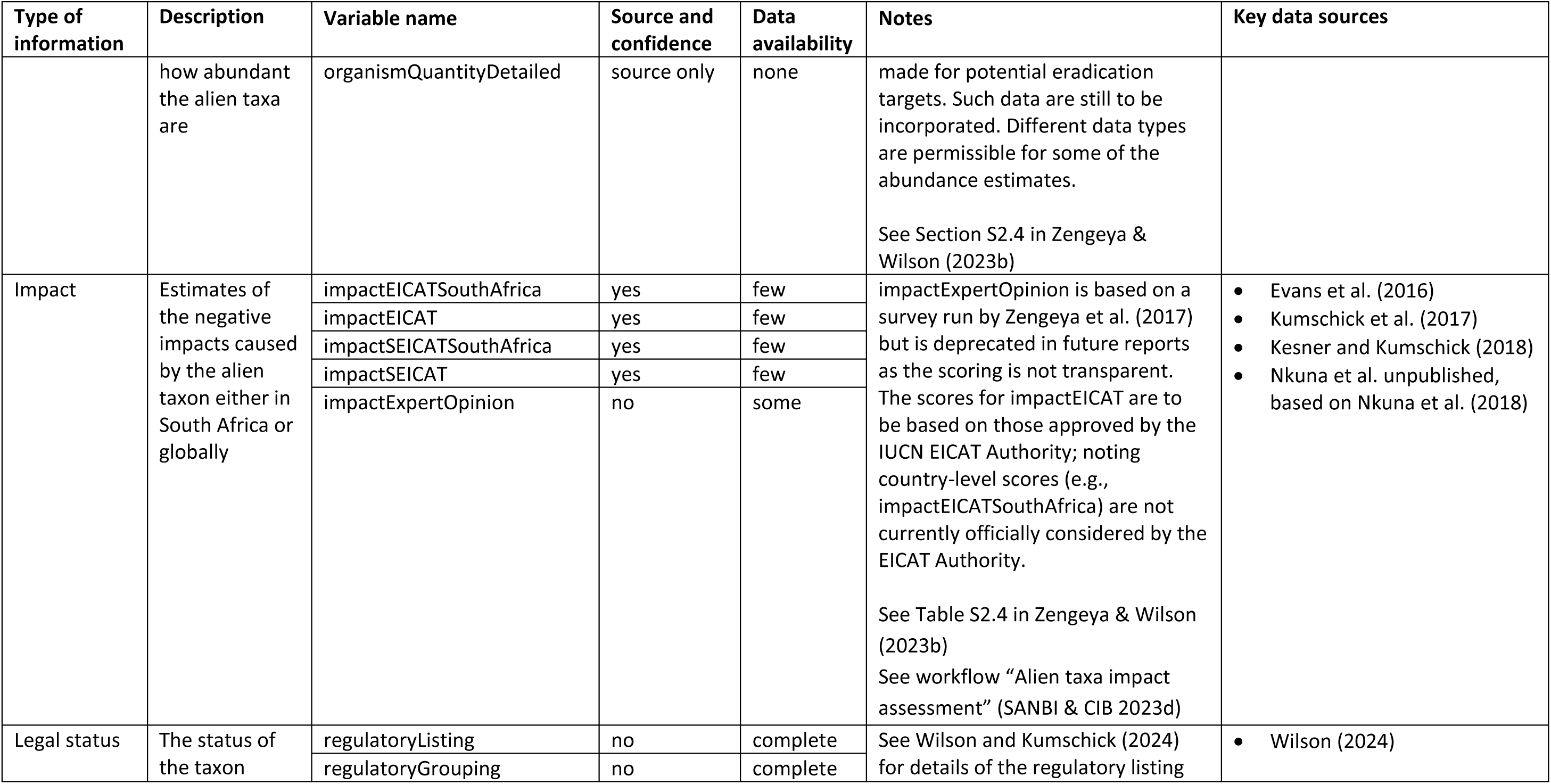
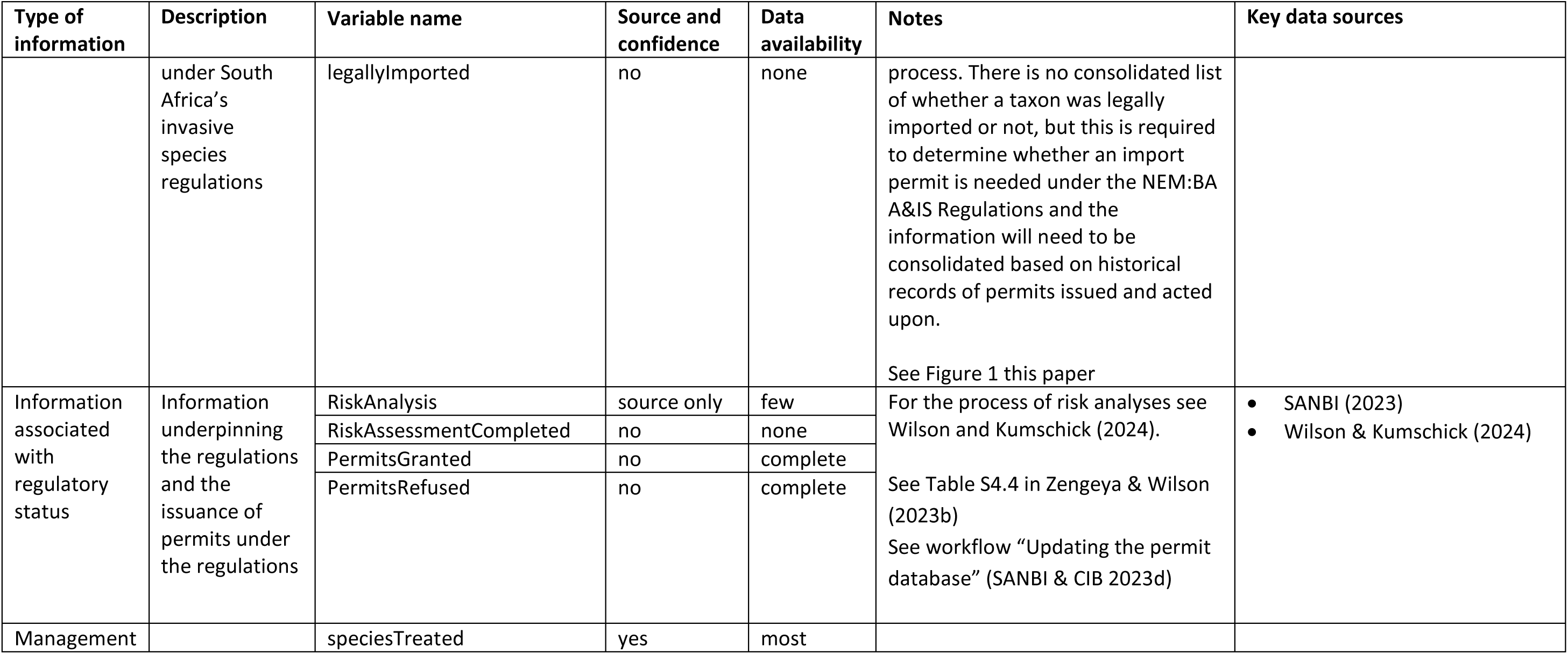
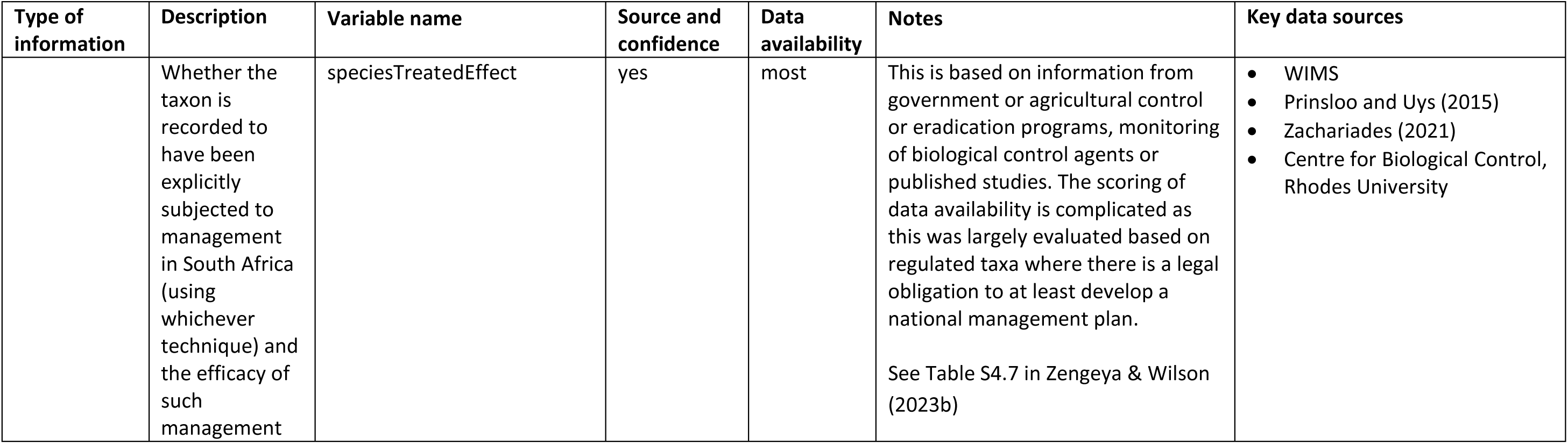
A summary of the metadata for the list of alien taxa compiled as part of South Africa’s national status report on biological invasions and key data sources used to inform each variable. Key data sources for each type of information were cited for at least 10% of taxa. Variable names (each corresponding to a column in the database) align with the Darwin Core where appropriate. In cases where source and confidence level are given, these data are captured in a separate column in the database. Data availability was scored on a qualitative scale—complete (98–100%); most (50–98%); some (5–50%); few (<5%); none—for those taxa for which the value was relevant [technically this excludes instances scored as not applicable (NA) and only considers instances scored as not evaluated (NE), though the distinction between NA and NE is not consistently captured in the current version of the database]. Data availability does not consider the level of confidence, in many cases this is low. Details of how the information informs the indicators used in the status report is in the full metadata. The list of alien taxa on the Prince Edward Islands (SANBI and CIB, 2023b) includes the same variables, although for some variables the factor levels differ. The full metadata are available in SANBI and CIB (2023c). The general workflow is shown in Figure 1, with further details and specific workflows in SANBI & CIB (2023d). Sources for data: Botanical Database of Southern Africa (BODATSA); Global Biodiversity Information Facility (GBIF); Plants of the World Online (POWO); 430 Southern African Plant Invaders Atlas (SAPIA); South African National Biodiversity Institute (SANBI); Water Information Management System 431 (WIMS).

### 2. Processing

An intermediary file was created to store the extracted, digitised information from a data source or from several related data sources, based on broad categories such as organism types (e.g., microbes, plants, freshwater fishes), habitats (e.g., marine taxa) or pathways (e.g., pet trade).

#### 2a. Standardisation

The data were systematically curated with metadata that provide details of what information is contained in each column, and what the different levels in each column mean (https://doi.org/10.5281/zenodo.8217211). The process to check taxonomic information was automated (see Faulkner, this issue) and is briefly described below. The nomenclature of non-plant taxa was checked against the Global Biodiversity Information Facility taxonomic backbone (GBIF; https://doi.org/10.15468/39omei). For plant taxa, the nomenclature was first checked against the Plants of Southern Africa database (NewPOSA; https://posa.sanbi.org) and if the information was not available, the Plants of the World Online database was used (POWO; https://powo.science.kew.org). However, for many taxa other user-defined taxonomic backbones were needed. For example, plant taxa not found in NewPOSA and POWO were checked against the International Plant Name Index (IPNI; https://www.ipni.org/) and names of animal taxa not found on GBIF were checked against Nemaplex (http://nemaplex.ucdavis.edu/) for nematodes, the World Register of Marine Species (WoRMS; https://www.marinespecies.org/) for marine taxa, and published inventories in other cases (e.g., for biological control agents released on plants, Zachariades 2021). Issues were flagged and noted in the intermediary file, such as unresolved terms, missing data and translated names (e.g., lumped or split names). Synonyms, names misapplied, names for which there were typos, or names with no authorship information provided in the original source were noted in the ‘otherNamesUsed’ column. The list of names in ‘otherNamesUsed’ is not intended to be exhaustive but is a pragmatic list so that people can find taxa that they otherwise might think are missing (i.e., if a name was used as the primary name for a taxon in at least one source it was included). A value of NA is possible for ‘scientificName’ for example for regulated species that are not valid taxa.

#### 2b. Status of alien taxa

The status of an alien taxon in South Africa was determined using information on its nativity, occurrence, degree of establishment, introduction status, and regulatory status (see below). The source was recorded and confidence estimated for all fields used to determine status, together with notes on any translations and interpretations.

##### isNative

isNative assesses whether the taxon’s native distribution range was within (at least a part of) South Africa. It is a factor with four levels (TRUE, FALSE, cryptogenic, data deficient). If a taxon is native to a part of South Africa (i.e., TRUE) then it does not belong in this dataset unless one of the following is true: the taxon has, or at some point had, native-alien populations (see Nelufule et al. 2022) in South Africa; the taxon is (or was) prohibited from being introduced to another part of South Africa under the NEM:BA A&IS List; a risk analysis (/assessment) has been conducted on the taxon for South Africa; and the taxon was at some point classified as alien to the whole of South Africa although the taxon’s nativity has since been settled and it is clear it is native. Cryptogenic taxa are of unknown biogeographic origin, and cannot be definitively categorised as native to any part of South Africa, but neither can be definitively categorised as alien to South Africa where they are present. Data deficient is when an assessment of biogeographic status is unfeasible because of uncertainty in the taxon identity.

##### occurrenceStatus

Evaluates whether a taxon occurs in South Africa (as of December 2022). It is a factor with four levels (absent, present, doubtful, and not evaluated). A taxon is noted as absent if an analysis of the available evidence suggests that the taxon is not present in South Africa or there is no evidence of presence. A taxon is assumed present if there is evidence to document the presence of the taxon in South Africa. The occurrence of a taxon is assumed to be doubtful if there is some evidence of the taxon having been present in South Africa, but there is doubt over the evidence or whether it is still present, including taxonomic or geographic imprecision in the records. The occurrence status of a taxon is noted as not evaluated if there has been no specific attempt to ascertain if the taxon is in (or has been in) South Africa.

Not all the values of the vocabulary for the Darwin Core term ‘dwc:occurrenceStatus’ are used (common, irregular, rare) as information on abundance is stored under ‘organismQuantity’. Values for ‘occurrenceStatus’ can inherit presences from ‘IntroductionStatus’ and ‘degreeOfEstablishment’ but can only inherit absences if used in combination with ‘isNative’ (e.g., native taxa that are not present outside of their native ranges, are still, of course, present).

##### degreeOfEstablishment

This variable specifies the degree to which the taxon, where it is alien in South Africa, is surviving, reproducing, and expanding its range. The coding is taken from the Unified Framework for Biological Invasions (Blackburn et al. 2011):

**A0-A1**: Never introduced beyond limits of native range to [a part of] South Africa (A0) OR was introduced but no longer present (A1)
**B1-C2**: Includes a range of taxa from those in quarantine to those that are reproducing outside of captivity or cultivation but where there is no clear evidence of having formed self-sustaining populations.
**C3-D1**: Taxa where there is naturalisation, and possibly spread, but there is no clear evidence of forming self-sustaining populations at a significant distance from point of introduction.
**D2-E**: Populations are self-sustaining a significant distance from the point of introduction

The wording and description are based largely on the Darwin Core term (dwc:degreeOfEstablishment; Groom et al. 2019), with the use of native rather than indigenous and the separation of A into A0 and A1 to indicate cases where taxa have disappeared from South Africa. In cases where a taxon can unequivocally be categorised at a certain level, but might be at a higher level, then the lower confirmed level should be used [e.g., if there is strong evidence of naturalisation (C3), but the evidence is ambiguous as to whether there has been spread (D1), or whether that spread resulted in new self-sustaining populations (D2), then that population(/taxon) would be scored as C3]. ‘degreeOfEstablishment’ can also be assessed as NA if ‘occurrenceStatus’ is absent or doubtful OR if ‘isNative’ is TRUE and ‘occurrenceStatus’ is present, and there is no indication of any individuals in what would be considered an alien range, or if it is not evaluated.

##### IntroductionStatus

This variable is a factor with three levels that provides a high-level classification of categories in the Unified Framework for Biological Invasions (Blackburn et al. 2011) that are used to assess ‘degreeOfEstablishment’: **A0-A1**: not currently present ‘NA’; **B1-C2**: introduced but not naturalised ‘presentAsAlienNotNaturalised’; **C3-D1**: naturalised but not invasive ‘NaturalisedNotInvasive’, and **D2-E**: invasive ‘Invasive’. It also specifically flags taxa with native-alien populations (sensu Nelufule et al. 2022). These are taxa that are native to a part of South Africa but have formed naturalised ‘NaturalisedNotInvasive:NativeAlienPopulations’ or invasive populations ‘Invasive:NativeAlienPopulations’ in another part of South Africa to which the taxon is alien. Native taxa with individuals in captivity or cultivation outside their native range in South Africa are not currently considered in this database.

##### Regulatory status

Regulatory status refers to whether a taxon was regulated as an invasive alien species under the NEM:BA A&IS Regulations (Wilson 2024). The values for regulatory status include: for listing taxa, the category of listing (‘1a’, ‘1b’, ‘2’, ‘3’, and ‘context-specific’); ‘Not.listed:was.listed’ for taxa that are not listed but were listed in the past; ‘Not.listed:was.proposed’ for taxa that were formally proposed for listing but never included on promulgated lists; ‘Uncertain’ for taxa where it is uncertain if they are listed or not (e.g., the identity of the taxon is at a higher level than the regulatory listing); and ‘Not.listed’ for taxa that are not currently listed. The listing is as per the NEM:BA A&IS Regulations of 2020. See Wilson and Kumschick (2024) for a review of how the lists have changed over time. Detailed notes on translated names and interpretations are available in the metadata (https://doi.org/10.5281/zenodo.8217211). If a taxon is not present in the country it is (or should be) regarded as unlisted, unless there is a discrepancy in the regulatory lists. For example, several freshwater crayfish species (*Faxonius limosus*, *F. rusticus*, and *Pacifastacus leniusculus*) are listed on the NEM:BA A&IS Regulations of 2020 but there is no evidence that they are present in South Africa.

#### 2c. Ancillary data

Additional fields in the intermediary file were used to capture information on whether the taxon was included in the species lists of previous reports, other information or motivations that needed to be flagged, and useful information on any other fields (pathways, distributions, impacts, and management) as per the species list if data are available.

### 3. Merging

Intermediary files were manually merged by standardised names and unification of ancillary data. Taxa with any flags on unresolved terms, missing data and translations were retained in the intermediary files and feedback provided to the source of the data for clarification and or correction. More than 10 intermediary files were created and to mitigate individual subjectivity, all databases were cross-checked against the checklist metadata by at least one member of the core author team (TZ, KF, JW).

**Literature**: The bibliography used is available in the checklist, a summary of key sources is presented in Table 1.

**Storage of data set:** online repository (http://dx.doi.org/10.5281/zenodo.14937470)

**License**: CC BY-NC 4.0.

**Format of data set**: digital xlsx file

**Version**: v1.1 http://dx.doi.org/10.5281/zenodo.14937470 (v1.0, http://dx.doi.org/10.5281/zenodo.8217197, was published along with the 2023 version of the national status report on biological invasions. v1.1 contains no new data but resolves some issues noted in v1.0, i.e., typographical and transcription errors and inconsistencies in the use of terms across variables).

**Language**: English

#### Data structure

Full metadata are presented on-line (SANBI and CIB, 2023c, http://dx.doi.org/10.5281/zenodo.7433113) with a summary presented in Table 1. The checklist has 85 fields that provide information on the status of alien taxa in South Africa, with information on their pathways, distributions, impacts, and management. The checklist attempts to adhere to the FAIR data principles. Specifically: data are Findable and Accessible through publication on the SANBI web-site (http://iasreport.sanbi.org.za) and an online repository (http://dx.doi.org/10.5281/zenodo.8217197); Interoperable by adapting fields to ensure, where possible, they conform to the Darwin Core data standards; and Reusable in terms of ensuring that it is easy to determine who generated the original data and obtaining permission for others to use the data. The associated data files were also produced in line with recommendations to make the data tidy—each row refers to a taxon and each column a particular variable with consistent units. The checklist also assesses the level of confidence (low, medium, high) for variables where there may be uncertainty following accepted best practice principles [see SANBI and CIB (2023a) for details].

### Summary of dataset

6 198 taxa were assessed for presence in South Africa of which there is evidence that 3 825 taxa are present, the presence of 1642 taxa is doubtful and 714 taxa are recorded as absent (Table 2). Over half of the taxa that are present are plants (2315 taxa), confirming the assertion that South Africa is a hotspot for plant invasions. Only a few of the taxa that are present were assessed as cryptogenic (28 taxa), and or native to some part of South Africa (257 taxa). Taxa that were assessed as absent were mostly animals (310 invertebrates and 165 vertebrates) and plants (211 taxa). Most of these taxa are either biocontrol agents that were released to control invasive plants but that did not establish, taxa listed as prohibited in at least one version of NEM:BA A&IS Lists, and or taxa included in previous status reports but there is no evidence that they are present. Doubtful taxa were mostly plants (1289 taxa) (presumed to be in cultivation, but there is uncertainty if they are present).

**Table 2.** The number and occurrence status of alien taxa in South Africa as of December 2022. Regulatory listing is as per the NEM:BA A&IS Regulations of 2020 and is grouped in two categories using the following descriptors: listed –taxa listed under various categories (‘1a’, ‘1b’, ‘2’, ‘3’, and ‘context-specific’); 2) Not listed - taxa that are not currently listed; and 3) Uncertain - for taxa where it is uncertain if they are listed or not (e.g., the identity of the taxon is at a higher level than the regulatory listing). Occurrence status was grouped into four categories using the following descriptors: 1) absent – a reasoned analysis of the evidence suggests the taxon is not present in South Africa; 2) present - there is evidence to document the presence of the taxon in South Africa as of December 2022; 3) doubtful - there is some evidence of the taxa having been present in South Africa, but there is doubt over the evidence or whether it is still present, including taxonomic or geographic imprecision in the records; and 4) not evaluated (NE) - there was no specific attempt, as part of this process for compiling the list, to ascertain if the taxon is in (or has been in) South Africa. Incertae sedis refers to taxa whose broad taxonomic relationships are unknown or undefined.

| Taxa | Regulatory listing | Occurrence status |  |  |  |
| --- | --- | --- | --- | --- | --- |
|  |  | absent | doubtful | present | NE |
| Animalia | Listed | 11 | 39 | 122 | 3 |
|  | Unlisted | 464 | 304 | 1263 | 1 |
|  | Uncertain |  |  | 2 |  |
| Bacteria | Listed |  |  |  |  |
|  | Not listed |  |  | 4 |  |
| Chromista | Listed | 3 |  | 1 |  |
|  | Not listed | 3 |  | 12 |  |
| Fungi | Listed |  | 2 | 2 |  |
|  | Not listed | 20 | 4 | 103 |  |
| Incertae sedis | Listed |  |  |  |  |
|  | Not listed | 1 |  |  |  |
| Plants | Listed |  | 17 | 366 | 13 |
|  | Not listed | 211 | 1276 | 1948 | 12 |
|  | Uncertain |  |  | 1 | 1 |
| Protozoa | Listed |  |  | 1 |  |
|  | Not listed | 1 |  |  |  |
| Total |  | 714 | 1642 | 3825 | 17 |

A fraction (13%) of the taxa that are present are listed under the NEM:BA A&IS Regulations lists of 2020. These are mostly plants (366 taxa) and animals (122 taxa). Some of the listed taxa were assessed as either absent (14 taxa) or their occurrence is doubtful (58 taxa). 1811 taxa were assessed for their introduction and establishment status in South Africa as of December 2022, and over a third of are invasive (719 taxa), 120 taxa are known to be naturalised but not invasive, and 329 taxa are present, but not naturalised (Figure 2). The status of remaining 4387 taxa is yet to be evaluated.

**Figure 2.**
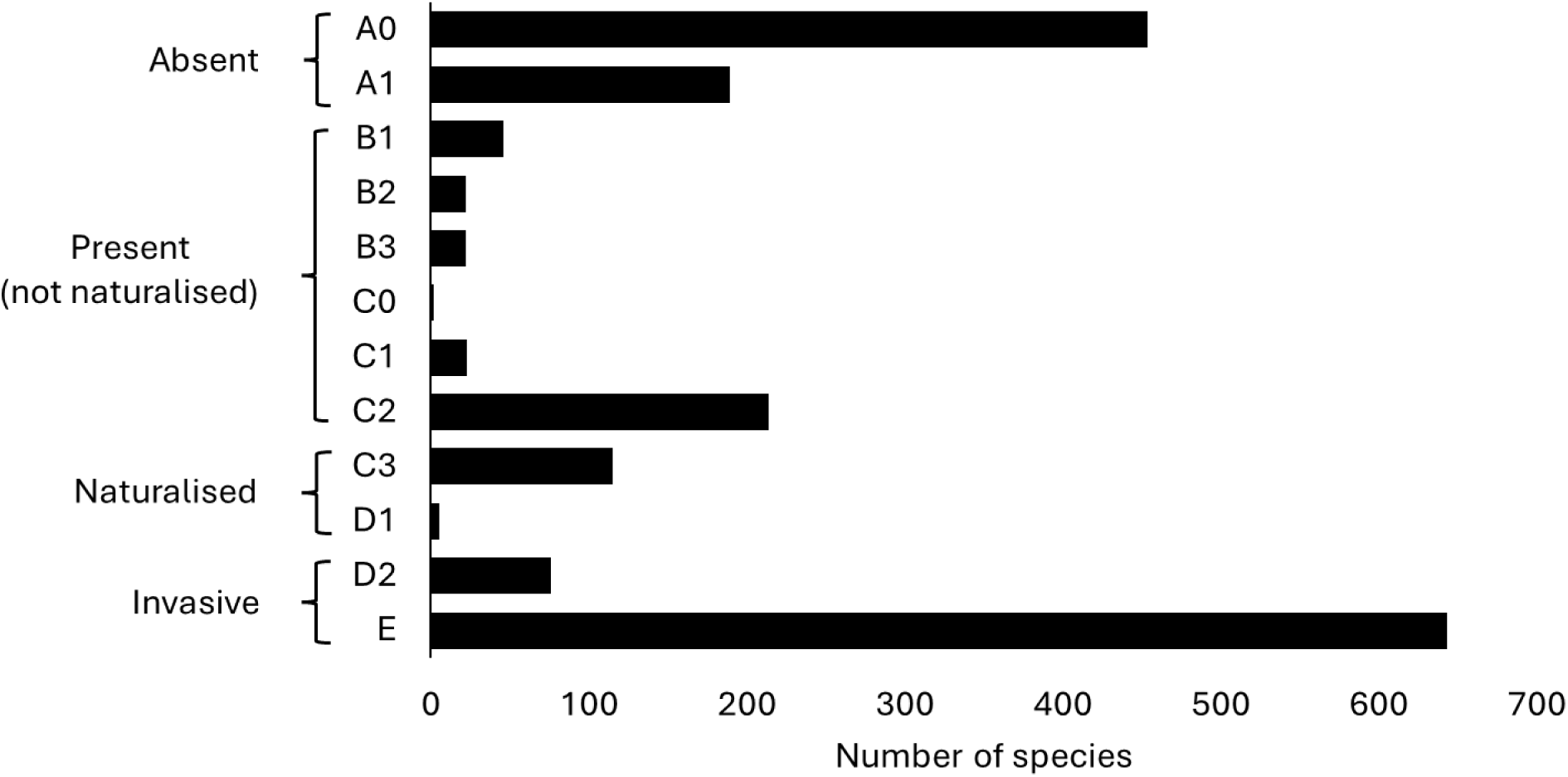
The introduction and establishment status of alien species in South Africa as of December 2022 as per the Unified Framework for Biological Invasions (Blackburn et al. 2011). Absent (A0-A1) – taxa that have never been introduced beyond limits of native range to [a part of] South Africa (A0) or were introduced but no longer present (A1); Present (not naturalised) (B1-C2) - includes a range of taxa from those in quarantine to those that are reproducing outside of captivity or cultivation but where there is no clear evidence of having formed self-sustaining populations; Naturalised (C3-D1) - taxa where there is naturalisation, and possibly spread, but there is no clear evidence of forming self-sustaining populations at a significant distance from point of introduction; and invasive (D2-E) – taxa with populations that are self-sustaining a significant distance from the point of introduction

### Limitations of dataset

Biological invasions are dynamic in nature, and there are often significant delays between when a taxon is introduced, when it is recorded, and when that record is reported and incorporated into a dataset. Therefore, this checklist, like all checklists of alien taxa, represents a snapshot of the situation. Many data sources have informed the checklist, but not all have been verified and integrated into the list. For example, due to a recent research focus, there are several lists of alien taxa in the pet and aquarium trades (e.g., Nelufule et al. 2020; Shivambu et al. 2022; Mantintsilili et al. 2022). However, there are many more alien taxa in captivity or under cultivation that are not yet included in the checklist, largely as data are not readily accessible or interoperable. Similarly, many groups and habitats have been under sampled. For example, information on most microorganisms is either not incorporated, is hard to incorporate as issues of nativity are unresolved, or there simply has been no sampling (e.g., few ecto-mycorrhizal fungi have been incorporated to date, despite their large conspicuous fruiting bodies; Magagula this issue). Thus, the checklist provides an underestimate of the number of alien taxa in the country, and this number will increase as new and historical data sources are verified and incorporated, and as new alien taxa are recorded and reported. As with many similar biodiversity datasets, the data that are generally available are on established or invasive alien taxa from large and charismatic groups.

The data sources that inform the checklist are highly variable as they were developed for various purposes, followed different methods, and implemented different standards and definitions. This checklist aims to provide a wide range of additional data on alien taxa in South Africa, however, not all the columns are presently complete. While in many cases this is because the primary research is yet to be performed (e.g., assessment of impacts), in others information exists that will be incorporated in future. These limitations mean that the checklist is not yet a complete baseline of the knowledge of which alien taxa are present in South Africa, nor of their status, pathways, distributions, and management. As such, differences between this list and the lists from previous reports (e.g., addition or removal of a taxon) need to be carefully interrogated, as they do not necessarily mean that there has been a change to biological invasions (e.g., increase or decrease in the number of alien taxa).

### Usage notes

The checklist can be used to report on the status of biological invasions in South Africa, track and report on progress towards Target 6 of the Kunming-Montreal Global Biodiversity Framework, and inform monitoring and management. The data can be integrated with other alien species checklists to obtain estimates at regional and global levels. However, the limitations detailed above mean that care needs to be taken when using the dataset to track trends. Change over time should be determined using a revised baseline that considers the recent incorporation of historical data, and when data were recorded, rather than when they were reported. We therefore encourage authors wishing to use this checklist to consult the corresponding author (TZ).

### Workflows

The overall workflow is presented in Figure 1, but specific workflows were developed to facilitate adding information to the checklist. In the third status report, seven workflows are outlined in SANBI and CIB (2023d). *Tracking data sources*, addresses process 1 on Figure 1. *Adding alien taxa and enrichment data to the species list*, addresses process 2 on Figure 1 including guidance for checking taxonomy and scoring nativity, occurrence status, degree of establishment (much of these details are précised in this paper), as well as incorporating information on pathways and introduction dates. *Alien taxa impact assessment* outlines a method to collate estimates of the negative impacts caused by the alien taxa either in South Africa or globally. *Updating the permit database* outlines how to incorporate particular information associated with regulatory status (permits issued or refused). *Money spent* outlines how to generate consolidated monetary estimates per taxon based on various sources. However, given the difficulties in disaggregating such data, these monetary estimates were not incorporated into the checklist. The final two (*Introduction pathway prominence* and *Sourcing, capturing, and reporting information for the Prince Edward Islands*) are not of direct relevance to the species list presented here.

## Discussion and Recommendations

The list presented here is the first consolidated list of all alien taxa in South Africa that aims at presenting data in a FAIR and tidy manner and that explicitly states the sources used and the confidence in the data presented. The list was constructed using set workflows (e.g., Faulkner this issue; Figure 1; SANBI and CIB, 2023d), and as such data have, as far as possible, been standardised and verified. However, while the list meets the mandated requirements to produce a list of invasive species under South African regulations (Department of Environment, Forestry and Fisheries, 2020), it is still not an appropriate baseline given key data sources have not yet been incorporated.

The status of alien taxa in South Africa, of course, changes over time. Taxa might need to be removed from the list due to taxonomic changes (e.g., a name becomes invalid), and their introduction status updated as more evidence of the current presence of taxon in South Africa become available (cf. Matthys et al. this issue). Taxa might be added either from existing data sources not yet incorporated or from new research and observations. A workflow has been developed to document such updates (*Tracking data sources*; SANBI and CIB, 2023d). Workflows are also needed to reduce the time between information being collected and when information is incorporated into the list, and ultimately to reduce the time between detection and action (Fernandez Winzer et al. this issue). More fundamentally, the list relies on the availability of data on biological invasions in South Africa. Capacity and resources for active and passive surveillance, the identification of new observations in the field, and the identification and classification of those observations are essential prerequisites if this list is to be an accurate representation of the alien taxa in South Africa. It is intended that this checklist will be updated and revised regularly as new information becomes available. Currently these updates are at least every three years, in line with the reporting requirements, but the intention is to move to annual updates, and aim for more frequent (quasi-real time) updates as soon as is practicable ensuring quality control processes are maintained.

We recognise that this list, as all such lists, contains errors. We are hopeful that by setting up a transparent process for creating the list, people will engage with what is here, provide feedback, and help correct errors. Please write to if you find issues. We similarly commit to providing feedback to data custodians when we identify issues while incorporating their information. In presenting these data we are thus presenting a structure and process that will enable us to move towards a dashboard while ensuring users can readily identify and report issues. By standardising access to data on biological invasions, we hope this checklist will help policymakers proactively address the issue (Zengeya et al. this issue; Groom et al. in prep).

## Supplementary material / links

For more information on the process used to compile the reports on the status of biological invasions and their management in South Africa see http://iasreport.sanbi.org.za

Copies of the latest report are also available at https://dx.doi.org/10.5281/zenodo.8217182 and associated appendices:

- Appendix 1. Species level pathway data http://dx.doi.org/10.5281/zenodo.8217192
- Appendix 2. The species list http://dx.doi.org/10.5281/zenodo.8217197
- Appendix 3. Metadata for the species list http://dx.doi.org/10.5281/zenodo.8217211
- Appendix 4. Workflows http://dx.doi.org/10.5281/zenodo.8217222
- Appendix 5. Pathways change tracker http://dx.doi.org/10.5281/zenodo.8217224
- Appendix 6. A database of permits issued under the NEM:BA A&IS Regulations 2014–2022 https://dx.doi.org/10.5281/zenodo.8229321

## Supporting information

Checklist of alien species in South Africa

## Authors credit statement

Authorship was based on those named as authors or contributing authors to the species chapter of the 3rd Report, and by evaluating authors and contributing authors to the other two reports based on them meeting two authorship criteria (e.g., data curation and methodology) before being invited to contribute further (e.g., review and editing).

Conceptualisation: BvW, JRUW, KF, TZ

Methodology: AS, BvW, EMcCJ, JRUW, KF, LFW, SK, SMT, WE, TZ

Data Curation: AS, BvW, EMcCJ, JRUW, KF, LFW, PM, TBR, SK, SMT, WE, TZ

Writing - Original Draft: JRUW, KF, PM, TZ

Writing - Review & Editing: All

## Acknowledgements

The lists are the result of many people who have worked on biological invasions in South Africa. We would particularly like to thank those who contributed as authors to the species chapters of the first and second reports: Andrew Turner, Charles Griffiths, Dai Herbert, Heather Terrapon, Ian Rushworth, Lee-Anne Botha, Lesley Henderson, Llewellyn Foxcroft, Michelle Greve, Musa Mlambo, Nonkazimulo Mdidimba, Pat Holmes, Pieter Winter, Rob Little, Tendamudzimu Munyai, Therese Forsyth, Trudy Paap, Tumelo Morapi, Xoliswa Ndeleni, Zanele Mnikathi; those who contributed information in the form of personal communications, in particular, Charlene Janion-Scheepers, Charles Griffiths, Costas Zachariades; and those who commented on drafts of the report when it was out for public review. Finally, we are very grateful to various people assisting with nomenclatural queries, in particular Caroline Mashau, David Richardson, Davina Saccaggi, Iain Paterson, John Bolton, Marieka Gryzenhout, Pieter Winter, and Simon Van Noort. We thank our late colleagues John Measey and Olaf Weyl for their contributions to the South African status reports and the science and management of biological invasions in South Africa. We thank the South African Department of Forestry, Fisheries and the Environment (DFFE) for funding, noting that this publication does not necessarily represent the views or opinions of DFFE or its employees. JW, KF, SK, PM, TZ also received funding from the B-Cubed project (Biodiversity Building Blocks for policy) which is funded by the European Union’s Horizon Europe Research and Innovation Programme (ID No 101059592). Views and opinions expressed are, however, those of the authors only and do not necessarily reflect those of the European Union or the European Commission. Neither the EU nor the EC can be held responsible for them.

